# Novel Machine Learning-based Approach to Identify Viral Biomarkers of Human Respiratory Emissions from Oral and Nasal Metagenomes

**DOI:** 10.1101/2025.09.26.678930

**Authors:** Kathryn Langenfeld, Peter Arts, Abigail Monahan, Allyson Criswell, Krista R. Wigginton, Melissa B. Duhaime

## Abstract

Humans spend approximately 90% of their lives in built environments, making virus transmission indoors a key determinant of health. Environmental sampling of respiratory viral pathogens is often challenging because of frequent non-detect measurements. Non-detect measurements do not differentiate between samples containing low or no pathogens from samples that simply lack respiratory expulsions altogether. This ambiguity can be resolved by scanning samples for a biomarker of human respiratory emissions. To do so, reliable biomarkers for environmental monitoring need to be identified. Ideal biomarkers are prevalent across individuals, abundant, and unique to the human respiratory tract. Here, we present a new machine learning-based approach to query for suitable biomarker candidates from publicly available metagenomes and apply it to identify viral biomarkers of healthy oral and nasal microbiomes. Twelve viral biomarker candidates were selected from 1,232 curated viral operational taxonomic units. The viral biomarker candidates had as much as 63% prevalence across respiratory metagenomes and prevalence was further increased to 77-81% by combining two or three biomarkers. Quantitative PCR confirmed that these viral biomarkers were prevalent and abundant in nasal swabs and saliva samples. Notably, top candidate biomarkers remained stable and detectable through multiple lab purification steps, increasing confidence in their viral origins and demonstrating their suitability for environmental monitoring. These findings demonstrate that existing metagenomes can be used to identify effective biomarker candidates for environmental sampling.

**IMPORTANCE:** Developing non-pharmaceutical interventions to reduce virus transmission indoors relies on robust environmental monitoring methods. Monitoring viral pathogens is challenging because of frequent non-detect measurements that introduce uncertainty. For instance, a non-detect measurement could indicate either the absence of the pathogen or simply the lack of human respiratory activity and thus exposure. To aid in distinguishing these scenarios, this study identifies viruses from the human respiratory tract using publicly available sequencing data that can be incorporated into environmental monitoring as biomarkers of human respiratory activity. These viral biomarkers will improve indoor monitoring to help enact interventions to mitigate virus transmission. Furthermore, our approach to identify biomarkers from existing metagenomes can be adapted for future biomarker identification in any system.

## INTRODUCTION

Environmental sampling of indoor spaces is critical to understanding respiratory pathogen transmission and developing effective nonpharmaceutical interventions, especially given that humans spend approximately 90% of their lives indoors (1). Respiratory viruses, such as influenza, coronaviruses, and respiratory syncytial virus, are spread indoors through transmission of droplets (2), aerosols (3), and fomites (4). A challenge of indoor air and surface sampling is the high frequency of non-detect measurements (5–7). Non-detect measurements can arise from multiple scenarios, including: (1) no pathogens were shed into the indoor space, (2) pathogens were shed into the space but were not detected due to issues with the analytical methods (e.g., below detection limit), or (3) pathogens were shed into the space but there were issues with the sample collection method (e.g., wrong surface sampled). Differentiating between these non-detect causes is important in transmission studies and in developing effective nonpharmaceutical interventions. To address this challenge, a biological indicator of human respiratory emissions, here termed a respiratory biomarker, is an ideal technical solution. With advances in shotgun sequencing, large repositories of publicly available sequencing data are currently untapped resources for the identification and development of potential biomarkers. A properly selected respiratory biomarker identified from publicly available sequence data could distinguish between pathogen non-detects measurements resulting from limited or no respiratory material in a sample and no pathogen shedding.

Biomarkers play an important role in public health and environmental monitoring by providing reliable indicators of human activity and exposure, thereby enabling accurate detection, assessment, and management of potential contaminants and disease transmission. Biomarkers have been widely implemented for monitoring fecal contamination and for interpreting wastewater epidemiology data. These are typically prokaryotes and viruses with high abundance and prevalence in human populations, such as coliforms (8), enterococci (8), crAssphage (9, 10), and Pepper Mild Mottle Virus (11, 12). For example, the United States Environmental Protection Agency bases its fecal pollution indicator policies on the concentration of coliforms and enterococci (13). In wastewater-based epidemiology studies, Pepper Mild Mottle Virus and crAssphage concentrations normalize to the proportion of fecal matter in wastewater by accounting for transient changes in population (14). However, these fecal biomarkers were selected without considering if they are found in other areas of humans such as on skin or in saliva. With advances in machine learning and viral bioinformatic tools, viral biomarkers can be identified that are prevalent and unique to particular environments. To date, biomarkers have not been identified for human respiratory emissions.

The most effective respiratory virus biomarker would be bacteriophages commonly found in the human respiratory tract, as their dissemination, persistence, and sampling recovery in air and surfaces would likely mirror those of viral pathogens. Many pathogens, including SARS-CoV-2 and influenza, have distinct tissue tropisms that results in different routes of shedding within the respiratory system (15). Therefore, it is necessary to identify biomarkers that represent different respiratory expulsion fluids, such as nasal mucus and saliva. Further, ideal viral biomarkers will be unique to the human respiratory tract and not found in other human samples, such as in stool or on skin. Lastly, as is the case for crAssphage and Pepper Mild Mottle Virus in stool samples, ideal viral respiratory biomarkers will be highly prevalent and abundant in humans, so they are able to be identified consistently in environmental samples.

Here, we developed an approach to identify viral biomarker candidates from existing healthy human metagenomes collected as part of the Human Microbiome Project (16). We created viral operational taxonomic units (vOTUs) from nasal and oral metagenomes, then applied various machine learning approaches that balanced biomarker candidate uniqueness to the human respiratory tract with prevalence and abundance in human oral and nasal metagenomes. Twelve viral biomarker candidates were then quantified in saliva and nasal swabs with quantitative PCR (qPCR) and compared to the abundance of three prevalent and abundant saliva bacteria populations previously proposed for use in forensics (17). These viral biomarkers can be applied in future environmental sampling studies and our approach may be utilized in the future to identify new biomarkers.

## MATERIALS AND METHODS

### Healthy human metagenome dataset download and quality control

A collection of healthy human metagenomes from the Human Microbiome Project (https://hmpdacc.org/hmp/) were downloaded from NCBI DbGaP (Project Accession phs000228, Table S1). A subset of available metagenomes was randomly selected using the sample_n function (dplyr, v1.1.4). Samples representing the respiratory tract included 108 anterior nares (i.e., nasal), 108 buccal mucosa, 8 saliva, and 19 throat metagenomes; non-respiratory tract samples included 97 stool and 61 retroauricular crease skin metagenomes (Figure 1A). Of the metagenomes selected, samples were collected from 164 healthy individuals with 48% (n = 78/164) of individuals identifying as female and ages ranging 18-40 years old. Quality control of reads was performed by trimming Illumina adaptors, then reads were decontaminated of PhiX174 with BBDuk (BBTools, v37.64). Bases with quality scores less than 10 were trimmed from reads, then trimmed reads with quality scores less than 10 or lengths less than 100 bp were removed with BBDuk. Human reads were removed using the Human Host Filtration Pipeline (18).

**Figure 1.**
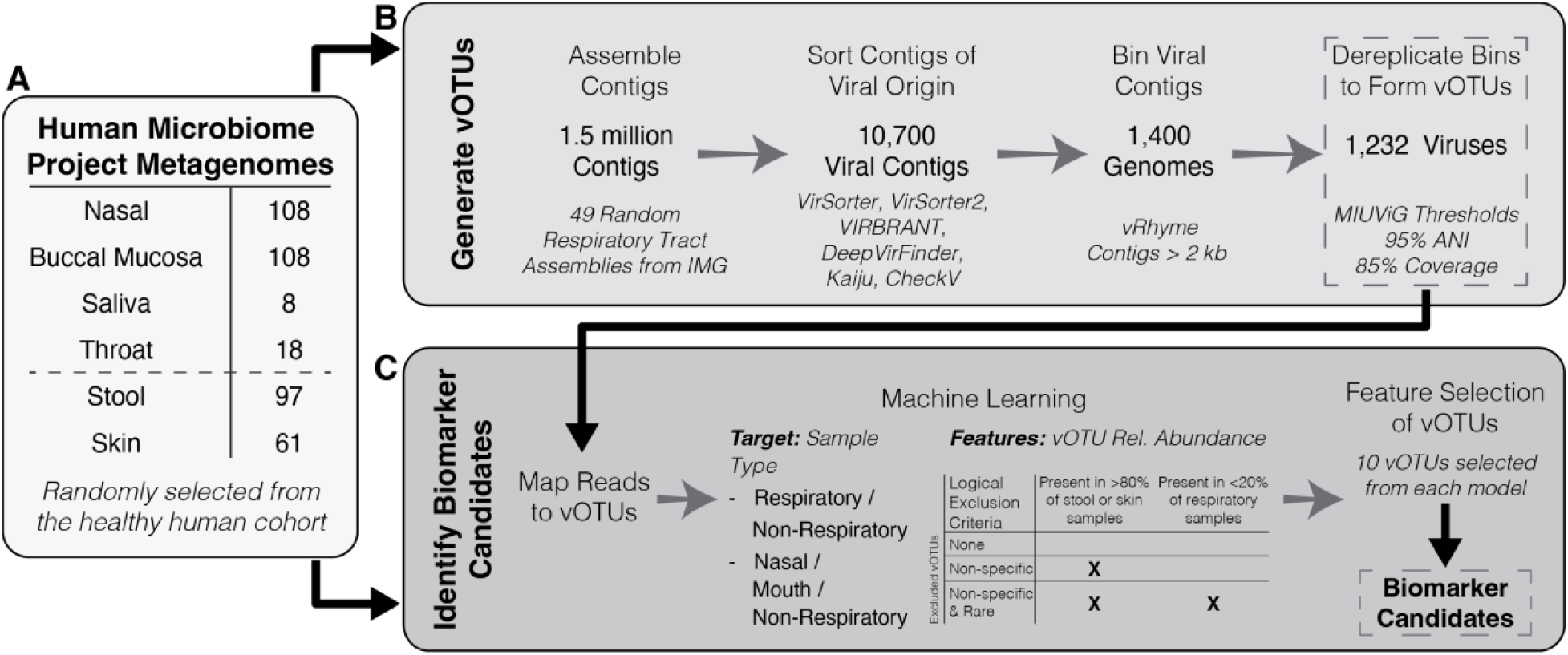
Summary of the process to identify viral biomarker candidates from healthy human metagenomes. (A) A random selection of metagenomes from the Human Microbiome Project was used to generate vOTUs and identify biomarker candidates. (B) A subset of the metagenomes had their assemblies downloaded from IMG to generate vOTUs. (C) Biomarker candidates were identified using machine learning modeling with the relative abundance of vOTUs across the metagenomes from the Human Microbiome Project.

### Generation of viral operational taxonomic units (vOTUs)

Assemblies from 49 random respiratory metagenomes (19 nasal, 27 buccal mucosa, 2 throat, and 1 saliva) were downloaded from IMG to curate viral operational taxonomic units (vOTUs) (Figure 1B, Table S2). To assess the likelihood of each contig being viral, six viral detection methods were run in sequence on contigs longer than 2,000 bp: VirSorter (v1.0.6) (19), VirSorter2 (v2.2.2) (20), VIBRANT (v1.2.1) (21), DeepVirFinder (v1.0) (22), Kaiju (v1.8.0) (23), and CheckV (v1.0.3) (24). Potential viral contigs were identified from these results using previously established rules (25). The viral contigs were binned using vRhyme (v1.1.0) (26) with a minimum contig length of 2,000 bp. The vRhyme bins labeled as “best” bins or circular viral contigs were retained. Viral bins from all of the samples were pooled and dereplicated with dRep (v3.4.3) (27) if bins shared greater than 95% average nucleotide identity and greater than 85% coverage of the shortest contig to form vOTUs (28). Viral sorting, binning, and dereplicating resulted in 1,232 vOTUs (Figure 1B).

### Read mapping to vOTUs, bacterial saliva biomarkers, and stool biomarkers

Read mapping was performed to assess the prevalence and uniqueness of vOTUs and other targets amongst the selected healthy human metagenomes (Figure 1C). We compared the prevalence and abundance of viral biomarker candidates to three commensal oral bacteria and a commonly used fecal biomarker, crAssphage. The three saliva bacteria and crAssphage are known to be highly prevalent and abundant in their respective matrices. The three saliva bacteria were *Streptococcus salivarius*, *Streptococcus sanguinis*, and *Neisseria subflava* (17). To map reads to saliva bacteria, the 3,000 bp region of their genomes including and surrounding previously developed PCR amplicon sequences were downloaded from NCBI (Table S3). The RefSeq crAss-like virus database was downloaded (08/24/2024) to have a reference set of dsDNA viruses commonly used as stool biomarkers (9). Often, a PCR primer assay, CPQ_056 (29), is used in environmental sampling that captures a subset of crAssphage (i.e., NCBI Accessions MW063138.1, MW067003.1, MW067002.1, MW067001.1, MW067000.1, MT006214.1, MK415410.1, MK415408.1, MK415404.1, MK415403.1, MK238400.1, NC_024711.1, BK049789.1, MZ130481.1, and MK415399.1). Five Bowtie2 (v2.4.2) indexes were built with default parameters for vOTUs, *N. subflava*, *S. salivarius*, *S. sanguinis*, and crAss-like virus targets. Read mapping with deinterleaved fastq formatted files of QC short reads was performed using Bowtie2 with the default mapping parameters. The number of basepairs mapping to each target was determined from Bowtie2 sam file output using flagstat (samtools, v1.11). The relative abundance of each target was calculated as the ratio of basepairs mapping to each target divided by basepairs in a sample.

### Biomarker candidate selection using machine learning

To identify vOTUs as candidates for human respiratory emission biomarkers, we used machine learning to select vOTUs that best distinguished respiratory samples from stool and skin samples (Figure 1C). We used supervised classification machine learning with kernelized support vector machines (Python v3.11) to handle our high dimensional data (30). In total, we created six different models by varying classification structure and logically excluding vOTU features. We utilized two classification structures: (1) respiratory (i.e., nasal, buccal mucosa, saliva, and throat) or non-respiratory (i.e., skin and stool) samples; and (2) nasal, oral (i.e., buccal mucosa, saliva, and throat), and non-respiratory samples. There were three sets of vOTU features used in our machine learning models: (1) all 1,232 vOTUs created; (2) removing non-specific vOTUs that excluded 105 vOTUs present in more than 80% of stool or skin samples; and (3) removing non-specific vOTUs and rare vOTUs that excluded 529 vOTUs present in more than 80% of stool or skin samples or present in less than 20% of respiratory samples. Models were created based on the relative abundances of vOTUs in each sample. The dataset was divided into training and test sets composed of 279 and 94 samples, respectively. Model accuracy measured the frequency that metagenome origin was correctly predicted. Ten vOTU features that best predicted sample origin classification were identified with forward feature selection. Model accuracy was measured again after feature selection. vOTUs selected as features by more than one model were chosen as viral biomarker candidates of human respiratory emissions. The biomarker candidates were combined to form “cocktails” to assess if having mixes of multiple targets would improve prevalence and abundance in respiratory samples. Every combination of two and three viral biomarker candidates was assessed for prevalence and abundance in respiratory and non-respiratory samples (Figure S1).

### Bioinformatic confirmation that biomarker candidates were of viral origin

To confirm that all of the selected biomarker candidates were of viral origin, the genomic content was assessed for each contig comprising the vOTUs (Table S4) and biomarkers were assessed in saliva and nasal mucus purified for viral particles. Two methods were used to assign taxonomy and functional potential. Taxonomy and functional potential were assigned with geNomad (v1.11 with database v1.9) (31) run end-to-end, which performs assignments by combining a neural network-based classifier and protein marker-based classifier. An alignment approach to taxonomic assignment was also run with megaBLASTn to viruses in the NCBI nucleotide database (performed on 3/19/2025). Functional annotation was also performed with Pharokka (v1.7.5) (32) using the PHROG reference database (performed on 8/3/2025) and PHANOTATE to predict genes along contig sequences. To establish confidence in viral provenance, we leveraged the knowledge that viruses have compact genomes with high coding density (33). The coding density of vOTUs was calculated as the total sequence length within open reading frames divided by the total sequence length. If taxonomy, functional potential annotation, and coding density resulted in uncertainty in whether the vOTU as a bonafide virus, further alignments were performed (on 4/1/2025) with BLASTn to the NCBI core nucleotide database, viruses in the NCBI Whole Genome Shotgun (WGS) contigs, and bacteria in the NCBI RefSeq Genome databases. Once the biomarker candidates were confirmed as viruses, the vOTUs were renamed based on where their prevalence was highest in respiratory environments (O = Oral; N = Nasal) and ranked based on prevalence and abundance in the metagenomic and qPCR (see details below) datasets.

### Laboratory confirmation that biomarker candidates were of viral origin

Viral particles were purified from pooled saliva and pooled nasal mucus to provide physical evidence that the selected biomarkers were of viral origin. Freshly collected 1-mL saliva and nasal swabs were collected from three individuals as approved by the Institutional Review Board (IRB-HSBS) at the University of Michigan (HUM00241431). The three saliva samples were pooled, then diluted with 7 mL of filter sterilized PBS with 0.5% bovine serum albumin and vortexed until homogenous. The nasal swabs were combined with 2 mL of PBS with 0.5% bovine serum albumin and vortexed for one minute. Initial 400-µL aliquots of the saliva and nasal mucus samples were collected and stored at 4°C until DNA extraction.

Size exclusion was performed on the remaining saliva and nasal mucus by filtering the saliva through a 0.22-µm PES syringe filter (Millipore Sigma, Cat. No. SLGPR33RS). 400-µL aliquots of filtered saliva and nasal mucus were collected and stored at 4°C until DNA extraction. Then, chloroform treatment was performed to lyse cells while not compromising most viral protein capsids with the exception of some enveloped viruses (34) and filamentous viruses (35). Chloroform treatment was performed by adding 500 µL and 100 µL of chloroform to the filtered saliva and nasal mucus, respectively, and vortexing for two minutes. The chloroform was removed from the sample by settling the sample on the benchtop for 15 minutes, then pipetting most of the chloroform from the bottom and aerating the trace remnants. Aeration significantly reduced the volume of saliva to 5 mL and nasal mucus to 300 µL. The nasal mucus was re-diluted to a volume of 800 µL using PBS with 0.5% bovine serum albumin for the remainder of the experiment. 400-µL aliquots of chloroform-treated saliva and nasal mucus were collected and stored at 4°C until extracting DNA.

Finally, non-encapsulated DNA was removed by adding 100 U/mL DNase I (Sigma Aldrich, Cat. No. 10104159001) that was suspended in a reaction buffer containing 750 µM Tris-HCl, 200 µM MgCl_2_, and 40 µM CaCl_2_. The DNase was allowed to react with the saliva and nasal mucus on the benchtop for one hour, then heat inactivated at 75°C for five minutes. Final 400-µL aliquots of DNase-treated saliva and nasal mucus were collected.

DNA extraction was performed in duplicate with 200 µL of all collected aliquots immediately after completing the experiment with a Kingfisher Flex instrument equipped with a 96-well attachment. Extractions were performed with the Applied Biosystems^TM^ MagMAX^TM^ Viral/Pathogen II Nucleic Acid Isolation Kit (Fisher Scientific Cat. No. A48383) with two wash cycles resulting in 50 µL elutions. qPCR was immediately performed on the DNA extracts.

### Saliva and nasal swab collection and DNA extraction

Saliva and nasal swabs were collected in the evening from ten individuals with no self-identified chronic respiratory diseases over the age of 18. Saliva samples were collected with SDNA-1000 saliva collection kits (Spectrum Solutions) and nasal swabs were collected with Quickvue Influenza Nasal Swab Tubes (Quidel) with 1 mL of PBS with 0.5% bovine serum albumin added. The study and all associated documents and protocols were approved by the IRB-HSBS at the University of Michigan (HUM00241431). Samples were stored at -80°C for a maximum of 36 days until DNA extraction. Duplicate extractions were performed on each sample. Immediately prior to DNA extraction, samples were thawed on ice. DNA was extracted from duplicate 200 µL saliva and 200 µL nasal PBS solution with a Kingfisher Flex instrument equipped with a 96-well attachment. The Applied Biosystems^TM^ MagMAX^TM^ Viral/Pathogen II Nucleic Acid Isolation Kit (Fisher Scientific Cat. No. A48383) with two wash cycles was performed with 50 µL elutions. DNA extracts were stored at -20°C for a maximum of one week until qPCR was performed.

### Biomarker candidate PCR primer design and qPCR reaction conditions

The abundance of viral biomarker candidates and bacterial saliva biomarkers were assessed in saliva and nasal swab DNA extracts using qPCR. Primers for the twelve vOTUs selected as viral biomarker candidates were designed using the IDT PrimerQuest^TM^ tool. The best primer assay for each vOTU was selected to maximize specificity. Specificity of each primer assay was assessed with NCBI Primer BLAST to the nr database where no or few hits indicated high specificity. Selected viral biomarker candidates’ primers and previously developed saliva bacterial biomarkers (17) primers are provided in Table S5. The qPCR threshold cycle (Ct) for each biomarker in the DNA extracts were compared to ddH_2_0 negative controls (NTC). qPCR reactions of duplicate saliva and nasal swab sample DNA extracts and duplicate virus purification pooled saliva and nasal mucus DNA extracts were performed with the QuantStudio 3 thermocycler (Thermo Fisher Scientific, Inc.). For each plate, one saliva and nasal sample were analyzed with two ddH_2_O negative controls for each primer set. The 20 µL reactions were prepared with 10 µL of Luna Universal qPCR mastermix (New England Biolabs, Cat. No. M3003), 0.5 µM of forward and reverse primers, and 5 µL of 1:10 diluted template. The reactions consisted of initial denaturation at 95°C for 60 seconds followed by 45 cycles of denaturation for 15 seconds at 95°C, then annealing and extension for 30 seconds at 60°C. The mean Ct value for duplicate reactions was divided by the mean Ct value of the duplicate NTC to determine the fold increase from NTC.

### Statistical analysis

All statistical analysis and figure creation was performed with R (v4.4.0) using ggplot2 (v3.5.1) and plotly (v4.10.4). Normality was tested with the Shapiro-Wilkes test. Wilcoxon’s test with Benjamini–Hochberg’s correction was performed on non-normal datasets.

### Data availability

Metagenomes were downloaded from the Human Microbiome Project (https://hmpdacc.org/hmp/) on NCBI DbGaP (Project Accession phs000228, Table S1). All assemblies were downloaded from IMG (Table S2). The specific contigs comprising the vOTUs identified by the machine learning models are listed in Table S4. Fasta files of the twelve vOTUs identified as biomarker candidates are available on Zenodo (10.5281/zenodo.17178845).

## RESULTS AND DISCUSSION

### Oral metagenomes generated more vOTUs than nasal metagenomes

To identify viruses pervasive in human respiratory emissions, we curated 1,232 vOTUs from 49 healthy human oral and nasal metagenomes (Figure 1B). The oral samples (i.e., buccal mucosa, saliva, and throat) had significantly more assembled viruses than nasal samples with oral and nasal assemblies having an average of 45 viral genomes and two viral genomes, respectively (*p*-value = 1.4x10^-7^; Table S2). A previous analysis of viruses in samples from the Human Microbiome Project identified significantly more vOTUs in oral than nasal samples (36). This observation may be due to fewer viruses comprising the nasal microbiome or issues with the sequencing data quality. In our study, sequencing depth after quality control and human read filtration was significantly greater in oral metagenomes than nasal metagenomes (*p*-value = 2.6x10^-5^). The oral assemblies also contained significantly more contigs longer than 1,000 bp than nasal assemblies (*p*-value = 4.9x10^-7^). Despite the nasal assemblies resulting in fewer contigs, the median contig lengths were not significantly different between oral and nasal assemblies (*p*-value = 0.26), indicating similar assembly quality between oral and nasal metagenomes. Previous reports comparing oral and nasal microbiomes are inconsistent regarding if the oral microbiome is more diverse than the nasal microbiome (37–40). Previous studies have found diverse viral communities in the oral cavity (36, 41–44). However, the limited prior work on viruses in nasal cavities resulted in few vOTUs suggesting low viral richness (36, 45). Future work should aim to further characterize the viral community and ecology of nasal cavities.

### 12 vOTUs were selected as viral biomarker candidates

Viral biomarker candidates of human respiratory emissions were identified from the vOTUs. Ideal viral biomarkers would be highly abundant and prevalent in respiratory samples and not found in other human environments, such as in stool or on skin. The identification of ideal candidates was based on: (1) identification of vOTUs that were indicators of oral, nasal, skin, or stool samples with machine learning approaches, (2) evaluation of the abundance and prevalence of vOTUs across target (i.e., oral and nasal) and non-target (i.e., stool and skin) body sites and the consistency of model vOTU selections, and (3) agreement across models as to the best choices of respiratory vOTU biomarker candidates.

Six machine learning models were applied to predict the origin of the 400 metagenomes based on vOTUs relative abundances across respiratory (i.e., oral and nasal) and non-respiratory (i.e. stool and skin) metagenomes. We performed modeling using three sets of features: all 1,232 vOTUs, removal of non-specific vOTUs (i.e., vOTUs present in more than 80% of stool or skin samples), and removal of non-specific vOTUs and rare vOTUs (i.e., vOTUs present in less than 20% of oral and nasal samples). The accuracy of the machine learning models ranged from 0.49-0.85 (Table S6), which indicated that our models were not overfitted to the training data. With this result, we proceeded to next apply feature selection to identify the ten vOTUs that best predicted sample origin from each model. The models were rerun with the selected ten vOTUs and the models’ accuracy range improved to 0.70-0.93 (Table S6). Across the six models, 40 vOTUs were selected by at least one model as key features (Figure 2A).

**Figure 2.**
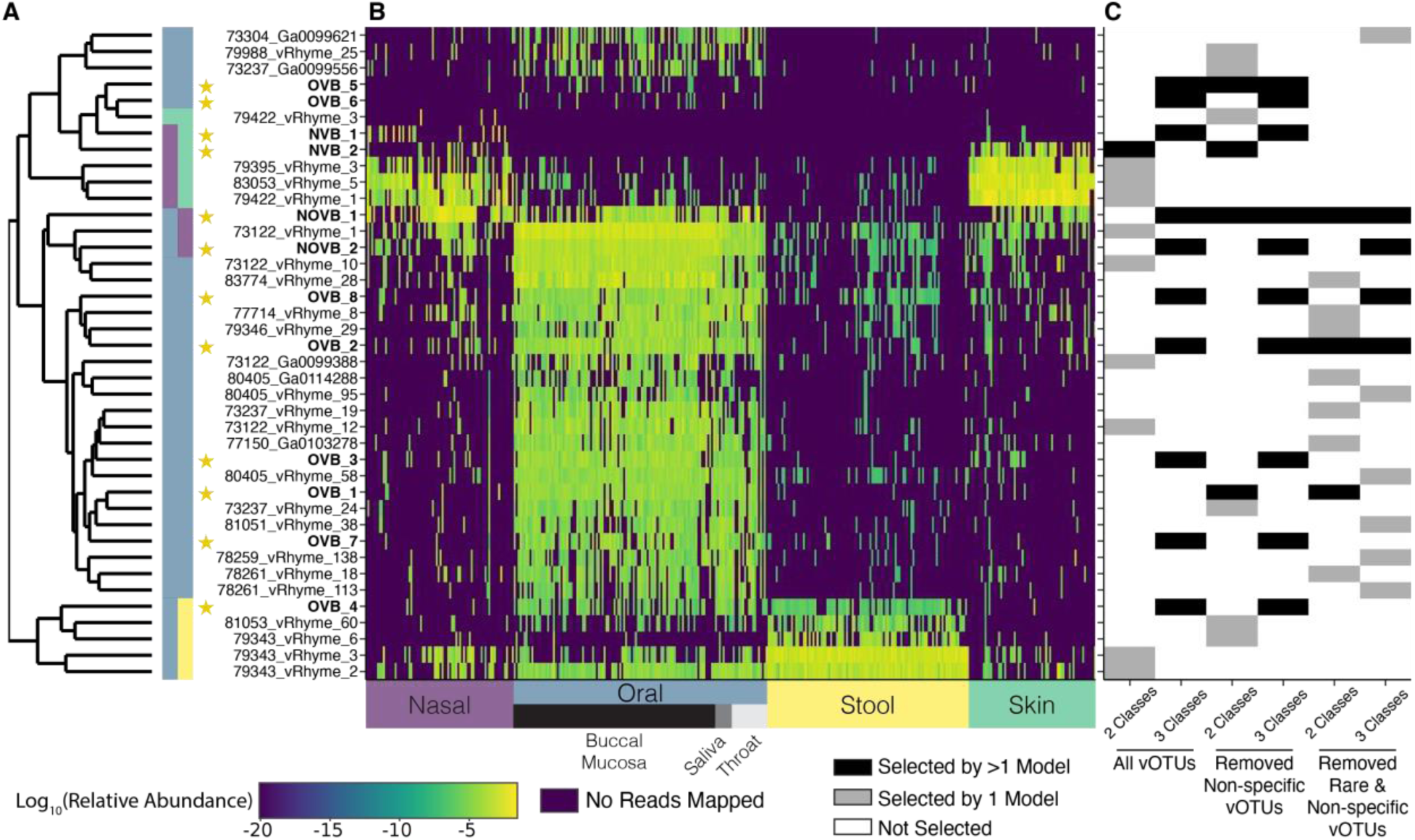
Relative abundance of vOTUs selected as key features in machine learning models across the HMP metagenomes. (A) vOTUs are organized by hierarchical clustering with euclidean distances based on relative abundances across all HMP metagenomes. The vertical color bar denotes clusters of vOTUs indicative of particular sample origins where nasal, oral, stool, and skin are represented by purple, blue, yellow, and green, respectively. The vOTUs identified by more than one model were selected as viral biomarker candidates, indicated with a star and bolded name. (B) Logarithmic relative abundance of vOTUs in each sample, where dark purple indicates a vOTU was not present in a Human Microbiome Project metagenome. The metagenome sample origin is indicated by the x-axis color bar. (C) Heatmap indicating which vOTUs were selected during feature selection in each of the kernelized support vector machine learning models. The models had two classification options to determine sample origins: two classes (respiratory or non-respiratory) and three classes (nasal, oral, or non-respiratory). There were three sets of features used: all 1,232 vOTUs, 105 non-specific vOTUs removed, and 529 rare or non-specific vOTUs removed (Figure 1C).

We next evaluated the abundance and prevalence of the 40 potential biomarkers in respiratory (i.e., nasal and oral cavities) and non-respiratory (i.e., stool and skin) locations to characterize which environments each vOTU inhabits. Of the 40 candidate vOTUs, 24 were highly abundant and prevalent in the oral environment exclusively (Figure 2A; Table S7). Although published studies on the viral component of these human microbiomes are sparse, previous research on bacterial communities has similarly shown that the oral microbiome is distinct from the other human microbiomes (37, 46). We identified three candidate vOTUs that were prevalent in nasal and oral samples (i.e., NOVB_1, 73122_vRhyme_1, NOVB_2; Figure 2A and B). Three vOTUs were highly unique to nasal (i.e., 79422_vRhyme_3) or oral samples (i.e., OVB_5 and OVB_6) but were not considered prevalent in any human environment (<15% prevalence in all sample types; Figure 2A and B). Ten of the candidate vOTUs were poor candidates for biomarkers of respiratory emissions given their high prevalence on skin or in stool: five vOTUs were prevalent in nasal and skin samples with five other candidate vOTUs prevalent and abundant in oral and stool samples (Figure 2A and B). Although these skin and stool vOTUs aided models in determining sample origin, they are not ideal respiratory biomarker candidates. The identification of these vOTUs prevalent in stool reinforced the decision to incorporate logical exclusion criteria in four of the models to exclude vOTUs that were highly prevalent in the non-target stool or skin samples (Figure 2C). The overlap of vOTUs in skin and nasal metagenomes aligns with previous findings of similar microbiota in skin and nasal samples (37). Some correspondence in the distribution of stool and oral vOTUs is not surprising, given prior observations of overlap between the oral and distal colon microbial communities (47) that has given rise to hypotheses that the oral microbiome may seed the gastrointestinal tract (48). Our data suggest this also may hold for human virome dispersal, as some vOTUs were prevalent and abundant in both oral and stool samples. However, more robust ecological analyses of the human virome are needed to evaluate these hypotheses, given that our trends are based on the analysis of only a few key vOTUs, rather than the thousands of vOTUs identified in the Human Microbiome Project data.

In our pursuit of candidate respiratory emission viral biomarkers, we next looked for agreement between the machine learning models. Across the six models, 12 vOTUs were selected as key features by more than one model (Figure 2C). The two most often selected vOTUs, NOVB_1 and OVB_2, were identified by five and four models, respectively. NOVB_1 and OVB_2 were commonly found in oral samples with NOVB_1 also found in nasal samples. The remaining vOTUs were selected by two (n = 7) or three (n = 3) models with greatest prevalence in oral samples (n = 4), nasal and skin samples (n = 2), nasal and oral samples (n = 1), or oral and stool samples (n = 1). Two vOTUs, OVB_5 and OVB_6, were highly specific to oral samples although were rarely present in the metagenomes. Given that these 12 vOTUs were identified by more than one model, all were further evaluated as viral biomarker candidates in this study.

### Biomarker candidate genomes demonstrate evidence of viral origins

We next evaluated whether the 12 biomarker candidates identified were of viral, rather than bacterial or human, origins. All contigs that comprised the 12 candidate vOTUs were confidently identified as being of viral origin based on a previously developed and rigorously evaluated viral contig sorting algorithm (25). To increase our confidence in this viral assignment, we sought additional evidence, including (1) viral taxonomy assignment, (2) identification of known viral proteins on the contigs, or (3) a high coding density characteristic of viral, rather than cellular, genomes. Based on our taxonomy assignment approach, all biomarker candidates were assigned as viruses by at least one method, except for NOVB_1 and OVB_1 (Table S4). Viral proteins, or those known to be associated with viral functions, were identified on the contigs of the biomarker candidates, with the exception of NOVB_1 (Table S4). The coding density of the vOTUs was high (90-98%; Table S4) compared to that expected of the human genome (1-2%) (49) for eleven of the biomarker candidates, with the exception of NOVB_1, which had a lower coding density of 71% (Table S4). These observations supported the hypothesis that these contigs were of non-human origin, given that viruses and bacteria have compact, efficient genomes with high coding densities (33, 50). With the exception of NOVB_1, all candidate biomarkers fit expectations of viral origins based on taxonomic assignment, gene annotation, and coding density.

To evaluate the persistence of viral biomarker candidates through various virus enrichment steps, we collected fresh saliva and nasal mucus samples from three individuals and subjected them to three sequential viral purification methods: size exclusion, chloroform lysis, and DNase degradation of non-encapsulated DNA. Initially, most biomarkers were present in at least one of the sample types, except for NVB_1 and OVB_8 (Table S8). Size exclusion tested if the viral biomarker candidates could pass through 0.22-µm pores, which is a typical size threshold for viral particles, with some exceptions for giant phages (51, 52). All of the biomarkers except OVB_7 were present after 0.22-µm filtration (Table S8), indicating that most of the biomarker candidates fall within the conventional viral size range. Chloroform treatment was then applied, which disrupts lipid-containing cell membranes, while leaving viral protein capsids intact—though a few virus types, such as enveloped viruses (34) and filamentous phages (35), can be chloroform-sensitive. After chloroform treatment, all remaining biomarker candidates were measurable in at least one of the samples, except for OVB_5 (Table S8). Next, DNase treatment was used to degrade any free DNA not protected by a viral protein capsid. Most of the remaining viral biomarker candidates were not measurable after this step, only NOVB_1, OVB_1, and OVB_4 remained in at least one sample (Table S8). The lack of detection of certain viral biomarker candidates after both chloroform and DNase treatment could be due to these candidates having capsids susceptible to chloroform, having most of their DNA free or in compromised capsids prior to DNase treatment, or low initial concentrations that fell below detection thresholds after the purification steps or dilution. Notably, NOVB_1 and OVB_1 were measurable after each purification step in both saliva and nasal mucus.

We further evaluated the contigs comprising the NOVB_1 vOTU, the only biomarker without any viral genes annotated, to determine whether the putative vOTU was misassigned as viral, or if instead it represented novel, not yet characterized viral diversity, referred to as “viral dark matter” (53, 54). Methods for identifying viral dark matter rely on deep learning methods, such as DeepVirFinder (55), and alignments to viral contigs from shotgun sequencing (56). DeepVirFinder indicated that both NOVB_1 contigs were of viral origin (*p*-values = 0.032 and 0.033). When NOVB_1 was aligned to the core nucleotide database (BLASTn), it aligned best to the *Homo sapiens* chromosome 5 (NCBI accession OZ171101.1; bit score = 10,761). However, we also searched for homology in the whole genome sequencing database (NCBI WGS) and identified that the NOVB_1 contigs aligned best to two assembled phage contigs (bit scores = 73.4 and 66.2). Based on this evidence, combined with its high coding density (71%) relative to that expected of the human genome (1-2%) (49) and that it was measurable after three viral purification steps, we concluded that NOVB_1 is a novel phage sequence that represents undescribed and uncharacterized sequence space of “viral dark matter”.

### Most viral biomarker candidates were more prevalent and abundant in respiratory samples than non-respiratory samples

We next evaluated the prevalence and abundance of the 12 vOTUs biomarker candidates across the metagenomes to simulate a quantitative survey across body sites. Most of the viral biomarker candidates had a greater prevalence in respiratory samples (nasal and oral) than non-respiratory samples (stool and skin) (n = 10/12; Figure 3; Table S9). Seven of the viral biomarkers were present in more than 40% of respiratory samples (range = 41-63%) and were less commonly found in non-respiratory samples (range = 6-39%). Of these biomarker candidates, five were specific to oral samples and two were found in both oral and nasal samples (Table S9). Both of the viral biomarkers not observed in oral samples were relatively rare (less than 10% prevalence) across respiratory samples. In total, four of the viral biomarkers (OVB_5, OVB_6, NVB_1, NVB_2) were present in less than 10% of respiratory samples (Figure 3) and were selected by the machine learning models without the rare vOTUs roughly excluded (Figure 2C; Table S9).

**Figure 3.**
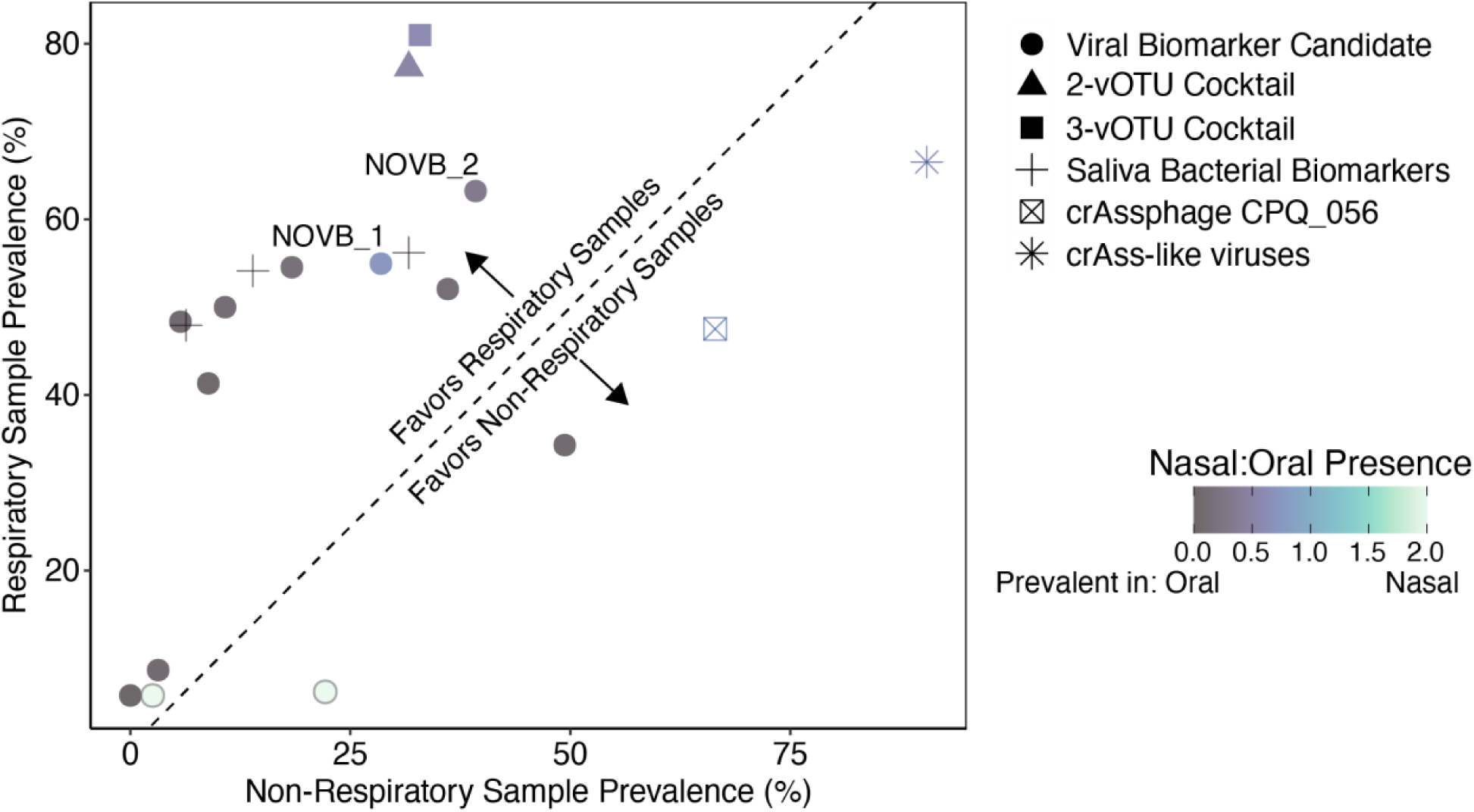
The prevalence of viral biomarker candidates, best viral biomarker candidate cocktails, three saliva bacteria from Jung et al. (2018), and crAssphage are compared across the different sample origins. The best viral biomarker candidate cocktails included a two-vOTU cocktail (NOVB_1 and OVB_1) and a three-vOTU cocktail (NOVB_1, OVB_1, and NVB_1). The points indicate the percent of samples with a target present in non-respiratory compared to respiratory samples. The dashed 1:1 line divides which targets are more prevalent in respiratory or non-respiratory samples. Points are shaped by viral biomarker candidates, two-vOTU cocktail, three-vOTU cocktail, saliva bacterial biomarkers, crAssphage CPQ_056 amplicon, and all crAss-like viruses as circles, triangles, squares, plus sign, crossed box, and asterisk, respectively. Points are colored based on the ratio of the prevalence in nasal compared to oral samples.

All of the viral biomarker candidates had a greater mean relative abundance in respiratory samples than non-respiratory samples (Figure 4; Figure S2A). For ten of the viral biomarkers, the mean relative abundance in respiratory samples was significantly greater than non-respiratory samples (*p*-values < 7x10^-4^). For two of the relatively rare viral biomarkers, OVB_5 and NVB_2, the mean relative abundance was not significantly greater in respiratory samples (*p*-values = 0.71 and 0.27, respectively). The best performing viral biomarker (in terms of abundance), NOVB_1, had the greatest mean relative abundance in oral (0.0106%; n = 84/108) and nasal (0.0585%; n = 49/108) samples (Figure 4; Figure S2A).

**Figure 4.**
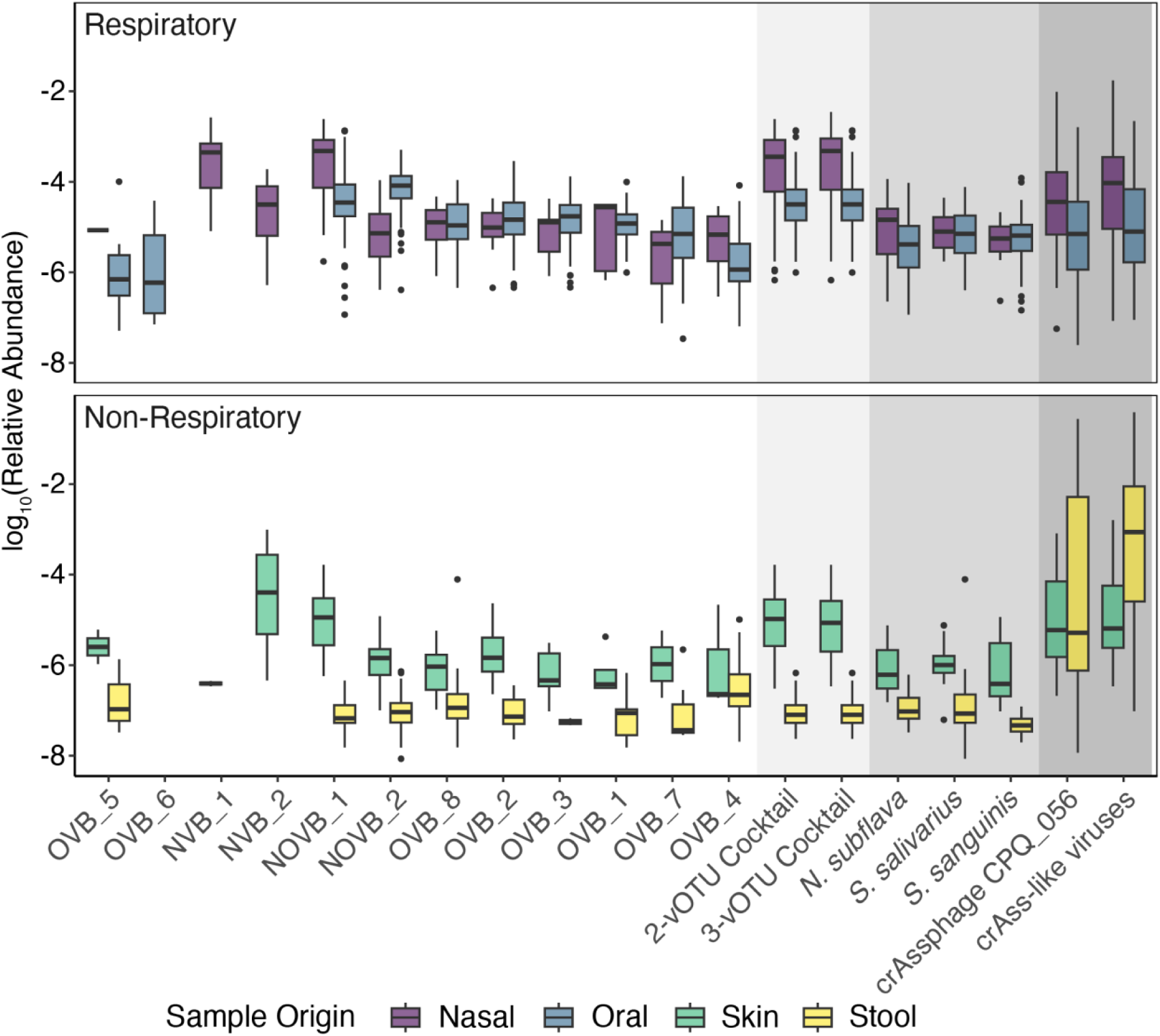
The abundance of viral biomarker candidates, best viral biomarker candidate cocktails, three saliva bacteria from Jung et al. (2018), and crAssphage are compared across the different HMP metagenome sample origins. The best viral biomarker candidate cocktails included a two-vOTU cocktail (NOVB_1 and OVB_1) and three-vOTU cocktail (NOVB_1, OVB_1, and NVB_1). Boxplot of logarithmic relative abundances for targets across the different sample origins where nasal, oral (buccal mucosa, saliva, and throat), skin, and stool are represented in purple, blue, green, and yellow, respectively. Any instances where a target was not present in a sample were not included.

### Viral biomarker candidates had similar prevalence and abundance to accepted standards

In order to benchmark our observations with accepted standards in the field, the prevalences of viral biomarker candidates were compared to three bacterial saliva biomarkers (17) and a commonly used viral fecal biomarker, crAssphage (9). The prevalence of the five viral biomarkers specific to oral samples was similar to that of the saliva bacteria biomarkers (Figure 3-4). CrAssphage was highly prevalent in respiratory and non-respiratory samples, with the prevalence of all crAssphage exceeding the most prevalent viral biomarker candidate, NOVB_2, in respiratory metagenomes (Figure 3). crAssphage CPQ_056 primers (29) and all crAss-like viruses were not specific to stool with high prevalence across all human microbiome samples. Previously, Tisza et al. (2021) found that most viruses are prevalent at only one body site with approximately 5% viruses prevalent in at least two sites, including some crAss-like viruses (36).

In addition to prevalence, we assessed if the viral biomarkers were ample in oral or nasal environments by comparing the observed relative abundances of viral biomarker candidates to the abundance of saliva bacteria biomarkers and crAssphage. The three bacteria species are known to be abundant in saliva (17, 37, 48) and crAssphage is the most abundant phage in stool (57). In comparison to saliva bacteria biomarkers and crAssphage in respiratory samples, the viral biomarkers were of similar relative abundances (Figure 4).

When combined, the saliva bacteria biomarkers were more prevalent in oral (n=133/134) than nasal (n=23/108) samples. However, when the saliva bacteria biomarkers were present, their mean relative abundances were similar in oral and nasal samples (0.00311% and 0.00207%, respectively). These results indicate that the saliva bacteria are abundant in both oral and nasal environments when present. Two of the viral biomarkers were significantly more abundant in nasal samples than the mean abundance of the three saliva bacteria biomarkers (*p*-values = 8.7x10^-9^ and 1.3x10^-5^ for NOVB_1 and NVB_1, respectively; Figure S2B) and the other nine viral biomarkers present in nasal samples were not statistically different in abundance. Whereas two of the viral biomarkers were significantly more abundant than the bacterial biomarkers in oral samples (*p*-values = 0.0032 and 5.5x10^-18^ for NOVB_1 and NOVB_2, respectively; Figure S2B). In summary, several viral biomarker candidates were highly abundant in either nasal or oral cavities, with NOVB_1 abundant in both environments, indicating its potential as a marker for respiratory monitoring.

CrAss-like viruses were highly prevalent and abundant in oral (n=102/134; 0.00992%) and nasal (n=59/108; 0.0742%) samples (Figure 4; Figure S2B and C). Only NOVB_1 was significantly more abundant than crAss-like viruses in oral samples (*p*-values = 2.1x10^-4^); however, NOVB_1 was significantly less abundant than crAss-like viruses in nasal samples (*p*-values = 0.0013). All other viral biomarker candidates were either significantly less than crAss-like viruses or similar in relative abundance in nasal and oral samples. CrAss-like viruses were previously shown to be present in multiple body sites (36). However, their high abundance may be partially due to their abundance comprising multiple populations (e.g., 268 crAssphage NCBI accessions) as opposed to a specific single vOTU, as is the case for the viral biomarker candidates.

### Grouping viral biomarkers into cocktails increases their prevalence in respiratory samples

While single viral biomarker candidates showed high prevalence and abundance, we explored the potential benefits of grouping multiple viral biomarker candidates into cocktail mixtures. In practice, these cocktails could be measured by performing separate assays for each biomarker using qPCR and then summing the results. Alternatively, a single assay could be conducted using a mixture of the primers, allowing the combined detection of multiple biomarkers within the qPCR reaction, thereby reducing reagent and supply costs. To facilitate this approach, we designed the primer sets for the viral biomarker candidates to have uniform reaction conditions and similar amplicon lengths.

We conducted an *in silico* assessment of all combinations of two-and three-vOTU cocktails (Figure S1). Based on their prevalence across all of the metagenomes, we identified a two-vOTU cocktail and a three-vOTU cocktail that showed high prevalence in respiratory samples (77.3% and 81.0%, respectively) and low prevalence in non-respiratory samples (31.6% and 32.9%, respectively; Figure 3). Both cocktails included NOVB_1, the most abundant biomarker in nasal and oral metagenomes, and OVB_1, a highly prevalent biomarker in oral metagenomes (Figure 2B). The three-vOTU cocktail also included NVB_1, which was unique to and highly abundant in the nasal cavity (Figure 2B). NOVB_1 alone was found in 62.7% of oral metagenomes, whereas the two-vOTU cocktail was found in 99.3% of oral metagenomes. In the nasal metagenomes, the three vOTUs cocktail was present in 58.3% of samples, compared to 45.5% for NOVB_1 alone. Both cocktails had minimal impact on prevalence in non-respiratory samples, with the two-vOTU and three-vOTU cocktail being present in 3.2% and 4.4% more of non-respiratory metagenomes, respectively. Although the cocktails increased prevalence in respiratory samples across individuals, the combined abundances of vOTUs in the cocktails was not significantly greater than NOVB_1 alone in any sample type (*p*-values = 0.53-0.94).

Overall, assessing multiple viral biomarker candidates increases the robustness of biomarker assays by reducing the likelihood of missing specific target populations and minimizing effects of sample-to-sample fluctuations. Due to variations in the human microbiome across individuals (47, 58), it is unlikely that everyone in a shared space over time will have a common vOTU in their respiratory fluids, especially considering our dataset was constrained to individuals from the United States. Others have demonstrated this challenge in fecal monitoring, whereby natural variations in fecal microbial composition between individuals impacts the utility of crAssphage and Pepper Mild Mottle Virus (PMMoV) biomarkers. For example, PMMoV is consistently shed across individuals, but its concentration in a single individual’s feces can be highly variable (12), plausibly due to fluctuations in diet (59, 60). On the other hand, crAssphage concentrations have been found to be stable within individuals, but can differ by six orders of magnitude between individuals (12). Such variations in PMMoV and crAssphage present challenges in environmental monitoring, particularly as community size decreases, because natural variations within and between individuals will result in large fluctuations in biomarker concentrations (14). Assessing multiple biomarkers with cocktail mixtures can reduce false negatives when assessing a sample for human respiratory emissions. By utilizing multiple biomarkers, natural variations in single vOTUs amongst and within individuals will not decrease the reliability of the assay as greatly.

### Viral biomarkers were found in saliva and nasal swab samples with qPCR

To further validate the efficacy of these viral biomarker candidates, we assessed their abundance in recently collected saliva and nasal swab samples with qPCR (Figure 5). Saliva bacteria biomarkers were measured alongside viral targets as a reference. In nasal swabs, any saliva bacteria biomarkers were found, at most, in half of the samples. There were five viral biomarker candidates that were more prevalent than the three saliva bacteria in nasal swabs (Figure 5A). Notably, NOVB_1 was found in all of the nasal swabs. In saliva, the bacterial biomarkers were highly prevalent, as previously reported (17). There were five viral biomarkers highly prevalent amongst saliva samples, with NOVB_1 found in all of the saliva samples.

**Figure 5.**
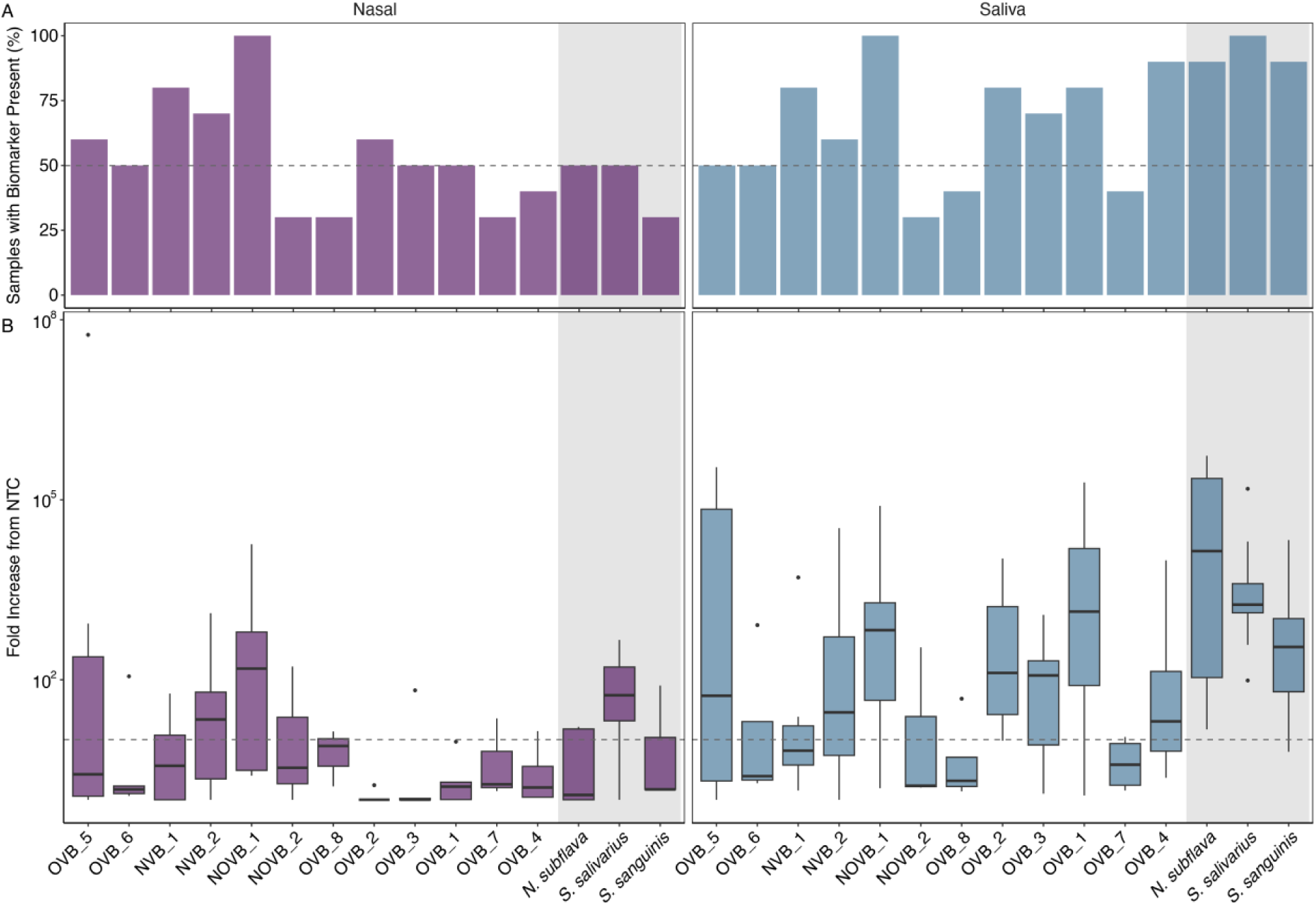
Presence and abundance of human respiratory emission biomarker candidates in nasal swab (n=10) and saliva (n=10) samples. (A) Percent of samples present for each biomarker candidate where present is defined as a greater than one-fold increase from the negative controls. The dashed line denotes when at least half of the samples had a biomarker present. (B) Boxplot of the fold-increase from the negative controls amongst the samples with the biomarker present. The dashed line indicates a 10-fold increase in biomarker abundance compared to the negative controls. The gray regions contain results for the bacterial saliva biomarkers from Jung et al. (2018).

The most abundant viral biomarker in nasal swabs, NOVB_1, aligned with the *in silico* observations from the Human Microbiome Project metagenomes. Only one other viral biomarker, NVB_2, had a mean fold increase above the NTC greater than an order of magnitude in nasal swabs. Among the saliva bacteria biomarkers, only *S. salivarius* had a mean fold increase greater than one order of magnitude above the NTC when present. Overall, the targets were more abundant in saliva samples, which may be attributed to lower overall biomass collected given the methodology. There were seven viral biomarkers with a mean fold increase from the NTC greater than an order of magnitude in saliva. The three saliva bacteria biomarkers were highly abundant in saliva with mean fold increases more than two orders of magnitude above the NTC. The viral biomarkers were generally less abundant with only two viral biomarkers, NOVB_1 and OVB_1, having mean fold increases greater than two orders of magnitude. In this study, we did not use probes for the saliva bacteria targets in our qPCR assays. This reduces the specificity of the assays based on NCBI primer BLAST results and may have inflated the qPCR abundances.

### Human respiratory viral biomarkers for environmental sampling

By assessing the prevalence and abundance of viruses in existing human metagenomes, we identified several viral biomarker candidates with high abundances and prevalences in newly collected nasal swabs and saliva. Three of the viral biomarkers, NVB_1, NVB_2, and NOVB_1, were highly prevalent in nasal and oral samples. There were four viral biomarkers, OVB_3, OVB_2, OVB_1, and OVB_4, that were prevalent and abundant in only oral samples. None of the viral biomarker candidates were unique to the nasal environment based on qPCR. The metagenomic results indicated that NVB_1 was unique to nasal samples; however, we observed an equally high prevalence in nasal swabs and saliva samples with qPCR.

Since pathogens are disseminated in air through droplets and aerosols (2, 3) generated from respiratory activities, pathogen-containing particles may originate from one of multiple locations in the respiratory tract, such as the oral cavity or nasopharynx. While influenza is shed from both saliva and nasal mucus, the route of expulsion can impact the transmissibility of the virus. The persistence of the virus may differ depending on the expulsion route and specifically the chemical composition of the matrix (61), and its transport in the environment is likely influenced by differences in particle sizes generated from different expulsion routes (62, 63). Therefore, identifying viral biomarkers unique to different particle origin sites would provide additional information on the persistence and transmission of pathogens in the environment.

Based on the qPCR results, NOVB_1 is the most promising viral respiratory biomarker candidate for environmental sampling because it is highly prevalent and abundant in saliva and nasal swabs. To increase the prevalence and abundance of biomarkers in nasal and saliva samples, multiple viral biomarkers could be combined to capture a wider population and increase the total abundance. Future work will assess the prevalence and abundance of these viral biomarker candidates on masks, surfaces, and air filters. Our results demonstrate the efficacy of implementing machine learning with viruses curated from existing metagenomes to identify biomarker candidates for environmental sampling. The methods developed here for biomarker identification can be utilized to improve other environmental surveillance applications.

## ACKNOWLEDGMENTS

This work was funded by Flu Lab as part of the Multidisciplinary InvesTIGAtion of Transmission to Ease inFLUenza (MITIGATE FLU) award. We thank the lead investigator of the MITIGATE FLU award, Linsey Marr, for her leadership of the research study and feedback from colleagues in the research study.

